# Prediction of prospective leaf morphology in lettuce based on intracellular chloroplast position

**DOI:** 10.1101/543538

**Authors:** Mio Hikawa, Kazuyo Nishizawa, Yutaka Kodama

## Abstract

In the accumulation response, chloroplasts move toward weak blue light, staying at positions along the periclinal cell wall. By contrast, in the avoidance response, chloroplasts move away from strong blue light, escaping to positions along the anticlinal cell wall. The accumulation response maximizes light capture and the avoidance response reduces photodamage. The intracellular positioning of chloroplasts is important for optimizing photosynthesis and may have common signals with the regulators that determine leaf morphology, another factor that affects photosynthesis. Here, we propose that intracellular chloroplast position can be used to predict prospective leaf morphology in lettuce (*Lactuca sativa*). To test this, we induced the accumulation or avoidance response in lettuce cells through exposure to the appropriate strength of blue light and observed the growth of the plants. Our results indicated that leaf area increased in response to weak blue light inducing the accumulation response, and leaf thickness increased in response to strong blue light inducing the avoidance response.

## 1. Introduction

Optimizing photosynthesis is important for regulating plant size and improving seed yield. For optimization of photosynthesis, the intracellular positioning of chloroplasts changes in response to environmental conditions such as light and temperature (Fujii and Kodama, 2018; Wada, 2013). For example, under strong light in warm conditions, chloroplasts localize along the anticlinal cell wall by moving away from the light, in a process termed the avoidance response (Wada, 2013). Similarly, under weak light in cold conditions, chloroplasts localize along the anticlinal cell wall, in a process termed the cold-avoidance response (Kodama et al., 2008; Ogasawara et al., 2013). The avoidance and cold-avoidance responses may reduce photodamage caused by chloroplast photoinhibition under warm and cold conditions, respectively (Fujii et al., 2017; Kasahara et al., 2002). In addition, under weak light in warm conditions, chloroplasts localize along the periclinal cell wall by moving towards the light, in a process termed the accumulation response. Physiologically, the accumulation response is believed to maximize photosynthesis, a theory that was recently confirmed experimentally through comparisons between various *Arabidopsis thaliana* mutants that show unusual chloroplast positioning (Gotoh et al., 2018). However, the effects of these responses in regard to the growth of wild-type plants from economically important species, such as vegetable crops, remained to be determined.

The chloroplast avoidance and accumulation responses are mediated by the blue-light (BL) photoreceptor phototropin (Jarillo et al., 2001; Kagawa et al., 2001; Sakai et al., 2001). Recently, we reported that phototropin is a thermosensory protein that mediates the cold-avoidance response (Fujii et al., 2017). Therefore, in many plant species, chloroplast positioning is altered by changes in BL intensity and in response to changes in temperature. Under artificial conditions such as those in a plant growth facility (or plant factory), chloroplast positioning should be influenced only by BL intensity, because temperature is normally kept constant. In this study, to investigate the effect of chloroplast positioning in plants under such conditions, such as vegetables grown in plant factories, we analyzed the effects on lettuce (*Lactuca sativa*) grown at a constant warm temperature and under strong and weak BL, which induce the avoidance and the accumulation responses, respectively.

## 2. Materials and methods

### 2.1. Plant materials and growth conditions

Lettuce (*Lactuca sativa*) seeds (No. 03503, Tohoku Seed Co. Ltd., Tochigi, Japan) were used. The seeds were planted in soil and cultivated under the tested light conditions. For growth test, the seedlings were grown under blue light-emitting diodes (LEDs) and red LEDs (ISL-150×150-H4RB, CCS Inc., Kyoto, Japan) lights in an incubator (IJ101, Yamato Scientific Co., Ltd., Tokyo, Japan) at 20°C for 21 days.

### 2.2. Observation of chloroplast positioning

For chloroplast observations, the seedlings were grown for 18 days under white light from fluorescent tubes in a growth cabinet (LH-240SP, Nippon Medical & Chemical Instruments Co., Ltd., Osaka, Japan) and then transferred to weak or strong BL (5 or 50 μmol m^−2^ s^−1^, respectively). To make it possible to clearly observe the mesophyll cells, a detached lettuce leaf was deaerated and permeated with water. Mesophyll cells were observed by light microscopy (BX60; Olympus, Tokyo, Japan) under a 100× oil-immersion objective (UPlanApo 100×/1.35 oil). The images were captured using a digital camera (DP72; Olympus) with cellSens software (Olympus).

### 2.3. Measurements of leaf area, thickness, and biomass

To measure leaf area, lettuce leaves were sandwiched in a clear plastic folder to flatten them, and images were captured using a scanner (imageRUNNER ADVANCE C5045, Canon Inc., Tokyo, Japan). From the images, leaf area was measured by ImageJ (Schneider et al., 2012), and the means and standard deviations were calculated. To measure leaf thickness, a digital caliper (100, MonotaRO Co., Ltd., Hyogo, Japan) was used, and the means and standard deviations were calculated. To measure biomass, the harvested aboveground biomass of the lettuce was measured as the fresh weight, and then oven-dried at 105°C for measurement of the dry weight. The means and standard deviations of the weights were calculated.

## 3. Results and discussion

To optically control chloroplast positions in lettuce cells, we used 5 μmol m^−2^ s^−1^ as weak BL (BL5) and 50 μmol m^−2^ s^−1^ as strong BL (BL50) and compared lettuce growth under the two conditions. When we irradiated lettuce cells with BL5 for 3 h, chloroplasts localized along the cell periphery at the top side of mesophyll cells and accumulated along the periclinal cell wall at the bottom side of the cell (Fig. 1A). This BL5-induced chloroplast positioning in lettuce cells is similar to that induced by weak light in plant species that grow under strong sunlight (Higa and Wada, 2016; Ishishita et al., 2016). When we irradiated lettuce cells with BL50 for 3 h, chloroplasts escaped from the periclinal position (Fig. 1B). In this study, BL5 and BL50 are referred to as the accumulation and the avoidance conditions, respectively.

**Fig. 1.**
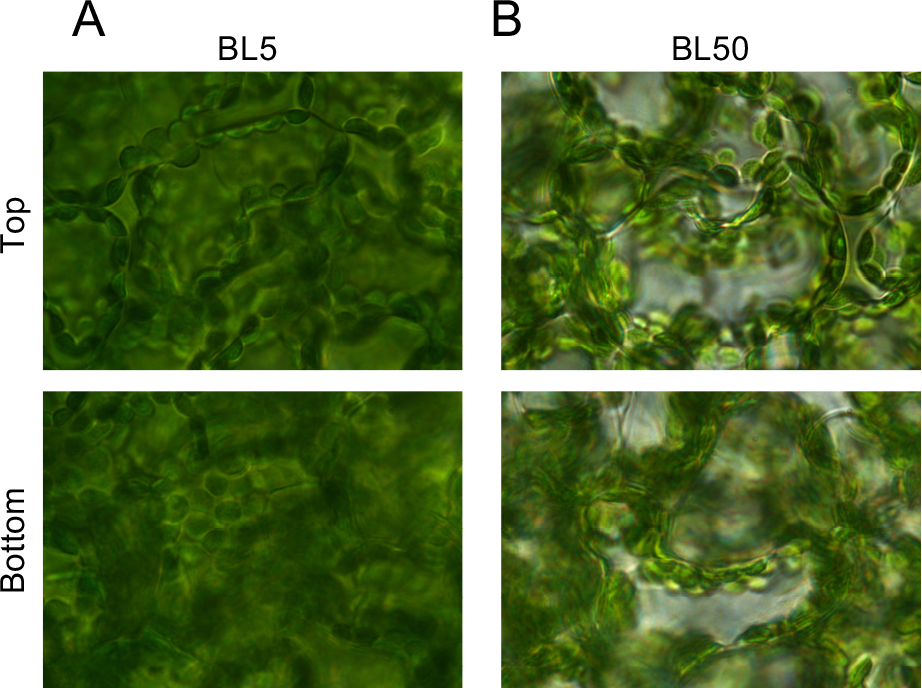
Optical control of chloroplast positioning in lettuce cells. (A, B) Intracellular chloroplast position in the accumulation response and the avoidance response in lettuce under BL5 (A) and BL50 (B), respectively.

For longer-term culture of lettuce plants, we added red light, 125 or 250 μmol m^−2^ s^−1^ (RL125 or RL250, respectively), to BL5 and BL50 to provide sufficient light energy for photosynthesis. Lettuce seedlings were incubated under the indicated BL and RL conditions at 20°C for 3 weeks. When combined with either RL125 or RL250, the BL5 seemed to enhance lettuce growth compared with BL50 (Fig. 2). To assess this quantitatively, we measured leaf area, thickness, and biomass. In combination with either RL125 or RL250, BL5 resulted in significantly larger leaf area compared with BL50 (Fig. 3A and B). However, the leaf thickness was significantly greater under BL50 than under BL5 (Fig. 3C and D). The results suggest that BL5 enhances leaf area and BL50 enhances leaf thickness. We then measured the fresh and dry weights of the aboveground biomass and found that fresh weight under BL5, in combination with either RL125 or RL250, was significantly higher than that under BL50 (Fig. 4A and B). Unexpectedly, however, dry weight under BL5 was comparable to that under BL50 (Fig. 4C and D), indicating that the biomass production was the same.

**Fig. 2.**
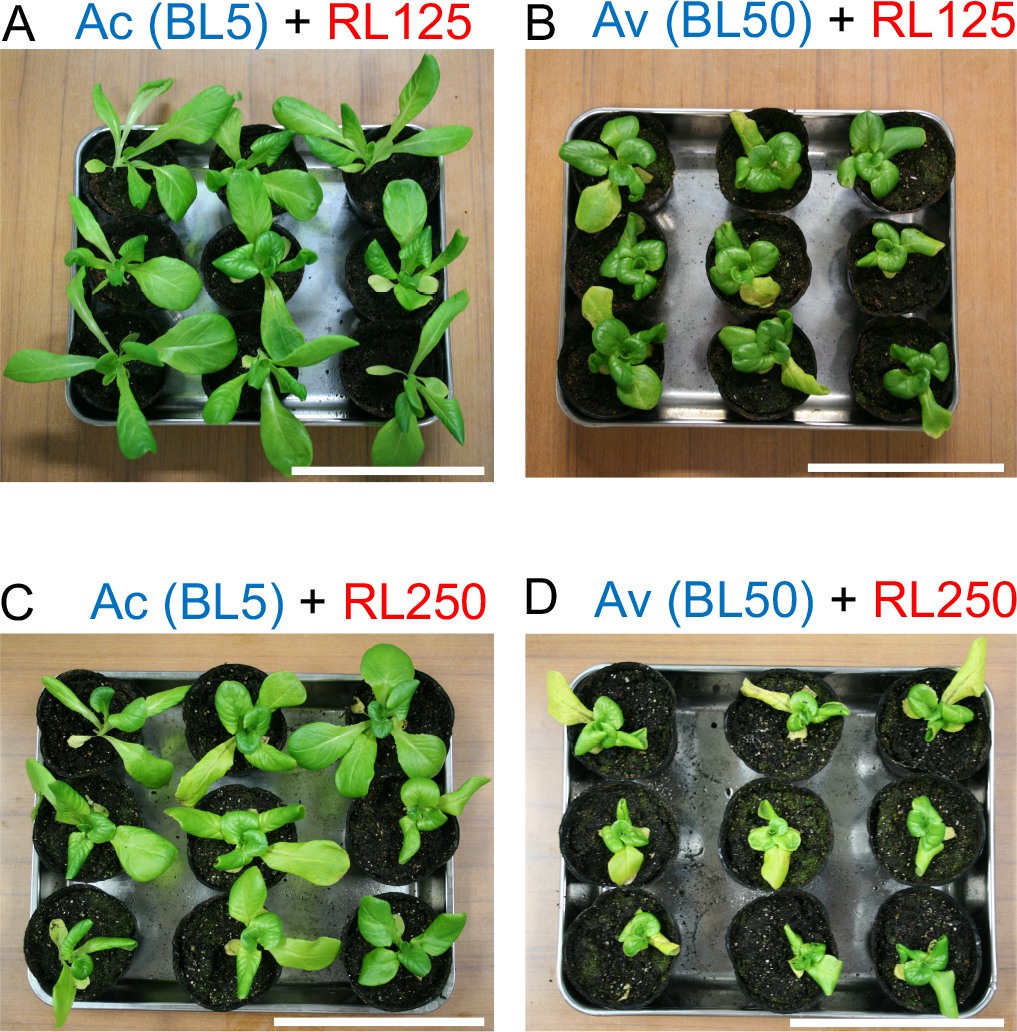
Lettuces cultivated under accumulation (BL5) and avoidance (BL50) conditions. (A, B) Photographs of lettuces cultivated under BL5 (A) or BL50 (B) in combination with RL125. (C, D) Photographs of lettuces cultivated under BL5 (C) or BL50 (D) in combination with RL250. White bars indicate 10 cm.

**Fig. 3.**
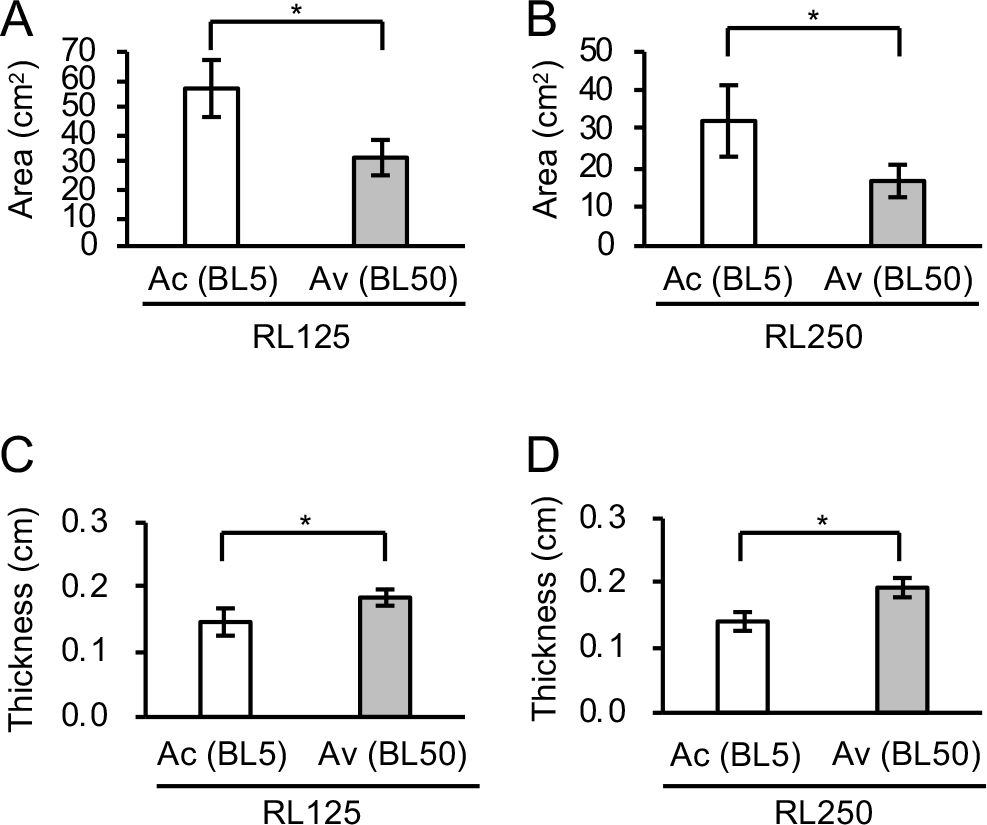
Leaf area and leaf thickness of lettuces cultivated under accumulation (BL5) and avoidance (BL50) conditions. (A, B) Comparison of leaf area in the BL5 and BL50 conditions in combination with RL125 (A) or RL250 (B). Bars indicates standard deviations. (C, D) Comparison of leaf thickness between the BL5 and BL50 conditions in combination with RL125 (C) or RL250 (D). In all panels, asterisks indicate statically significant differences between conditions (Student’s *t* test, *P* < 0.01).

**Fig. 4.**
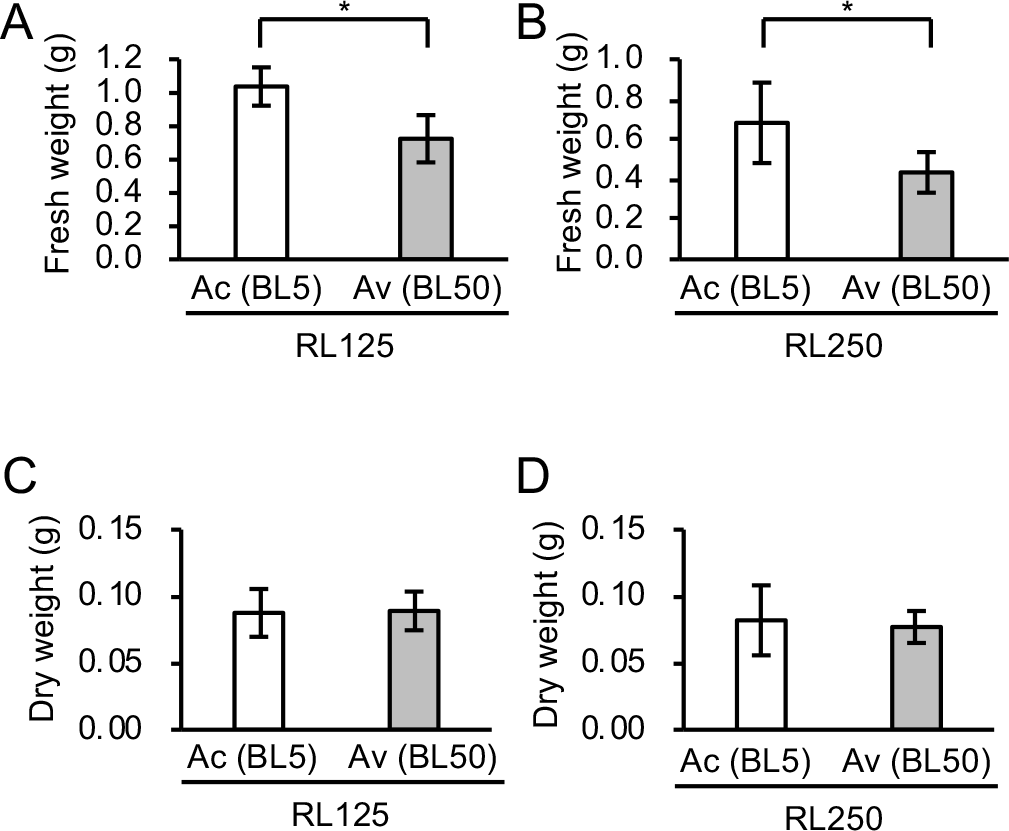
Fresh and dry weights of lettuces cultivated under the accumulation (BL5) and avoidance (BL50) conditions. (A, B) Comparison of fresh weights of leaves between the BL5 and BL50 conditions in combination with RL125 (A) or RL250 (B). Asterisks indicate statically significant differences (Student’s *t*-test, *P* < 0.01). (C, D) Comparison of dry weight between the BL5 and BL50 conditions in combination with RL125 (C) and RL250 (D). Student’s *t* test indicated no significant difference between the two samples (*P* = 0.86 in C and *P* = 0.63 in D).

In a previous study using *A. thaliana*, leaf area and biomass (fresh and dry weights) increased in conditions that induced the accumulation response (Gotoh et al., 2018). In the present study with lettuce, however, we found that the dry weight, and hence the biomass, was comparable under the accumulation (BL5) and avoidance (BL50) conditions (Fig. 4A–D), which is partially inconsistent with that report (Gotoh et al., 2018). It may be relevant that in our experiments we compared two physiological conditions (BL5 and BL50), but Gotoh et al. compared genetically distinct groups (wild-type plants and plants with mutations related to chloroplast positioning) (Gotoh et al., 2018). Further study will be needed to determine whether and how this difference in experimental design might have caused the inconsistency.

When we compared the fresh weights of plants grown under the two BL conditions, the weight for the plants grown under BL5 was much higher than that for the plants grown under BL50 (Fig. 4A and B). Given that the dry weights under the two conditions were comparable (Fig. 4C and D), this implied that lettuce grown under BL5 had a higher water content. Because higher BL intensity increases the degree of stomatal opening (Kinoshita et al., 2001), we suspected that the BL50 condition might result in more open stomata, resulting in greater transpiration from stomata and thus lower water content.

Although the leaf morphology was very different between the accumulation (BL5) and avoidance (BL50) conditions, the dry weights were comparable (Fig. 4C and D). Plant photosynthetic production under the two conditions thus seems to be same, suggesting that some form of compensatory growth occurs in lettuces under BL5 as compared to the BL50 conditions (namely, homeostasis). This compensatory growth may be based on varying strategies for leaf light capture during lettuce growth. In the BL5 accumulation conditions, because chloroplasts localize along the periclinal cell wall of the uppermost palisade cells, light penetration into deeper cells within the leaf could be expected to be inhibited (Gotoh et al., 2018). To efficiently capture light so as to maximize photosynthesis during growth, leaf area might be increased, or possibly leaf thickness might be decreased. By contrast, in the BL50 avoidance conditions, chloroplasts localize along the anticlinal cell wall of the uppermost cells. Because the light can penetrate into cells deeper in the leaf (Gotoh et al., 2018), leaf thickness might be increased to efficiently capture light, or leaf area might be decreased. Lettuce may thus have a regulatory mechanism to compensate for variations in chloroplast positioning.

Based on our observations, we propose that chloroplast position responds to a signal that also controls leaf morphology of lettuce. The relationship could also be direct—that is, the accumulation and the avoidance responses could serve as signals to enhance leaf area and leaf thickness, respectively. Based on these observations, we can predict prospective leaf morphology of lettuce by observing chloroplast position and we can modulate leaf morphology by altering light conditions. Given that chloroplast position can be changed by BL in many plant species, and that leaf morphology is important for many vegetable food crops, this control strategy may be employed for many economically important plants in artificial growth conditions.

## Acknowledgments

The authors thank for Dr. Eiji Gotoh (Kyushu University) for critical reading of the manuscript. The work was supported by the JST-A-Step (Y.K.) and the JST-ERATO (JPMJER1602: the Numata Organelle Reaction Cluster) (Y.K.).

